# The dynamic evolution of *Drosophila innubila* Nudivirus

**DOI:** 10.1101/197293

**Authors:** Tom Hill, Robert L. Unckless

## Abstract

Viruses coevolve with their hosts to overcome host resistance and gain the upper hand in the evolutionary arms race. *Drosophila innubila* nudivirus (DiNV) is a double stranded DNA virus, closely related to *Oryctes rhinoceros* nudivirus (OrNV) and Kallithea virus. DiNV is the first DNA virus found to naturally infect *Drosophila* and therefore has the potential to be developed as a model for DNA virus immune defense and host/virus coevolution within its well-studied host system. Here we sequence and annotate the genome of DiNV and identify signatures of adaptation, revealing clues for genes involved in host-parasite coevolution. The genome is 155555bp long and contains 107 coding open reading frames (ORFs) and a wealth of AT-rich simple sequence repeats. While synteny is highly conserved between DiNV and Kallithea virus, it drops off rapidly as sequences become more divergent, consistent with rampant rearrangements across nudiviruses. Overall, we show that evolution of DiNV is likely due to adaptation of few genes coupled with high gene turnover.

****Highlights**:**

- We sequence the genome of DiNV.
- Few genes are rapidly evolving between nudiviruses.
- We find high gene turnover between DiNV and its closest relatives.
- Helicase and ODV-E56 show consistent signatures of adaptation.

**Abbreviations**
DiNV: *Drosophila innubila* Nudivirus
OrNV: *Oryctes rhinoceros* Nudivirus
AcMNPV: *Autographa californica* Multiple Nucleopolyhedrovirus
ORF: Open reading frame
ODV-E56: Occulsion derived virus envelope protein 56
SNP: Single nucleotide polymorphism

## 1. Introduction

Baculoviruses and nudiviruses are large double stranded DNA viruses (90-180kbp genomes, 30- 300nm virions) that infect a wide array of arthropods (Jehle et al., 2006). They contain between 90 and 180 genes, of which a common set of 20 are key to the activity of the virus. Baculoviruses can usually be characterized by their helically symmetrical, rod-shaped nucleocapsids contained in stable occlusion bodies (known as polyhedra) and a viral encoded RNA polymerase (Jehle et al., 2006; Rohrmann, 2013). These factors allow the viruses to remain stable and infectious in most environmental conditions, and to remain active independent of the host RNA polymerase. Nudiviruses are close relatives of baculoviruses and while they are like other baculoviruses in many ways, they differ in the viral particle shape and that some do not form a baculovirus-like occlusion body (Wang et al., 2006). Currently there very few described nudiviruses and most infect arthropods, including fruit flies (*Drosophila*), rhinoceros beetles (*oryctes rhinoceros*), crane flies (*tipulidae*) and tiger prawns (*penaeus*) (Burand, 1998; Unckless, 2011; Wang et al., 2012). A subgroup of bracoviruses are also found within the nudivirus clade. These viruses are symbiotic with their host braconid wasp, making up a component of the parasitoid wasps venom (Bézier et al., 2009).

Though baculoviruses are among the best studied insect DNA viruses, we have limited understanding of how the arthropod immune system has evolved to suppress DNA viruses, and how the viruses in turn have evolved to escape this suppression. Recently, a nudivirus was discovered in the mushroom-feeding drosophilid species, *Drosophila innubila* (Unckless, 2011). The *Drosophila innubila* nudivirus (DiNV) is actually found across a large range of *Drosophila* species in the new world, found in every species collected in the Americas, varying in frequencies from 3% to ∼60% (Unckless, 2011). DiNV has been shown to reduce the viability of infected flies (infected flies survive 8-17 days post-infection, versus 20-31 days survival in mock infected controls), had significantly shorter lifespans in wild collected flies (median survival of 18 days and 43 days in virus infected and uninfected wild flies respectively) (Unckless, 2011). Infected females also laid significantly fewer eggs than uninfected flies (a median of ∼82% fewer offspring than mock infected controls). While not yet characterized in DiNV, other nudiviruses cause swollen, translucent larvae and increased larval deaths in their hosts (Burand, 1998; Payne, 1974). DiNV, like other nudiviruses, is suspected to infect the gut of infected adults and larvae. With the recently discovered Kallithea virus (Webster et al., 2015), DiNV has the potential to be developed into a powerful tool to study host-DNA virus interactions (Unckless, 2011) because of the wealth of resources available for studying the *Drosophila* innate immune system (Hales et al., 2015; Hoffmann, 2003).

To begin to gain an understanding of the host/virus coevolutionary arms race, we must start with a detailed characterization of the virus itself, including the sequencing, annotation and analysis of the viral genome. Here we sequence the DNA of an individual *D. innubila* male fly infected with DiNV and use the resulting metagenomics data to report the assembly and annotation of the DiNV genome. As found previously, DiNV is closely related to OrNV and the more recently found Kallithea virus. We find adaptive evolution across the genes in DiNV that is consistent with divergence based analyses across other baculoviruses and a population of *Autographa californica* Multiple Nucleopolyhedrovirus (AcMNPV) (Hill and Unckless, 2017). These results suggest that a few genes are adaptively evolving across this diverse group of viruses and that DiNV may be a useful model for understanding the evolution of a pathogenic DNA virus and the corresponding evolution of the host immune system.

## 2. Methods

### 2.1. Genome sequencing

Wild *Drosophila innubila* were captured at the Southwest Research Station in the Chiricahua Mountains between September 8^th^ and 15^th^, 2016. Baits consisted of store-bought white button mushrooms (*Agaricus bisporus*) placed in large piles about 30cm in diameter. A sweep net was used to collect the flies over the baits. Flies were sorted by sex and species at the University of Arizona and males were frozen at -80 degrees C before being shipped on dry ice to Lawrence, KS. All *D. innubila* males were homogenized in 50 microliters of viral buffer (a media meant to preserve viral particles, taken from (Nanda et al., 2008)) and half of the homogenate was used to extract DNA using the Qiagen Gentra Puregene Tissue kit (#158689, Germantown, Maryland, USA). We determined whether flies were infected by PCR screening for two viral genes, *P47* and *LEF*-4 (Supplemental Table 1 for primers and PCR conditions). The amplicons from flies screening positive for DiNV were sequenced (ACGT, Inc., Wheeling, IL, USA) to confirm the identity of the PCR product. One infected individual (ICH01M) was selected for sequencing. We constructed a genomic DNA library consisting of virus, *Drosophila* and other microbial DNA using a modified version of the Nextera DNA Library Prep kit (#FC-121-1031, Illumina, Inc., San Diego, CA, USA) meant to conserve reagents (Baym et al., 2015). We sequenced the library on one-twentieth of an Illumina HiSeq 2500 System Rapid-Run to generate 14873460 paired-end 150 base-pair reads (available at NCBI accession number SAMN07638923 [to be released upon acceptance of the manuscript]).

### 2.2. DiNV genome assembly

We used an iterative approach to assemble the DiNV genome. First, we trimmed all Illumina paired-end short reads using sickle (parameters: minimum length = 20, minimum quality = 20) (Joshi and Fass, 2011) and checked our data for any biases, high levels of PCR duplicates or any over represented sequences using FastQC. Ruling out these problems, we then mapped all Illumina paired-end short reads of the infected *D. innubila* fly ICH01M to a draft *D. innubila* genome (Robert L. Unckless, unpublished) using BWA MEM (parameters: -M) (Li and Durbin, 2009). Second, we took all unmapped reads and assembled them using Spades (default parameters) (Bankevich et al., 2012). Following this, we identified each contig’s closest hit via a BLASTN search to the non-redundant database with an E-value cutoff of 0.0001 (Altschul et al., 1990). Third, we took all contigs, including those with BLAST hits to any nudivirus or baculoviruses, and concatenated these to the draft *D. innubila* genome. We then re-mapped all reads to a preliminary *Drosophila innubila* genome, with the putative DiNV contigs attached (BWA mem parameters: -M) and collected all unmapped reads, as well as all reads mapping to the nudivirus or baculovirus contigs. We performed a further assembly using Spades with these reads, and assigning all nudivirus or baculovirus contigs as trusted contigs and all other previously assembled contigs with non-viral hits as untrusted (--trusted_contigs – untrusted_contigs). Finally, we repeated this process one further time, which yielded a 157429bp contig with considerable homology to nudiviruses. This contig has a mean coverage of 1124, a maximum coverage of 1887 and minimum of 116.

### 2.3. DiNV validation

We compared our assembled sequence with all known nudiviruses using MAFFT to identify aligned regions (MAFFT parameters: --auto) and its divergence from each other nudivirus (Katoh et al., 2002). We also remapped our short-read data to the *Drosophila innubila* genome with the viral genome concatenated (BWA mem -M) (Li and Durbin, 2009) and visualized it using IGV to identify any inconsistencies that may come with assembling a circular genome (Robinson et al., 2011), including the collapsing of duplicated regions, repeats of genes from the 'start' of the sequence onto the ‘end’ of the genome, or large structural rearrangements. While we found no large structural problems or duplication issues, we found inconsistent coverage across the last 1561bp of the sequence. This region showed strong homology to *Serratia liquifaciens*. The median coverage of the genome is 1124, while the median coverage of this *Serratia* portion is 157, suggesting either a misassembly or low frequency insertion.

We used pindel (default parameters) to attempt to identify further structural errors in our genome, but only confirmed our low confidence with the *Serratia* portion by its high frequency deletion (Ye et al., 2009). We concluded this region was not part of the consensus sequence due to its low coverage versus the rest of the genome and its low frequency found with pindel (0.128), though it may be a segregating horizontal gene transfer. To finally confirm or reject the presence of this *Serratia* portion, we designed primers across the edge of the *Serratia* portion and across the start/end of the DiNV sequence, labelled A-F in Supplementary Table 1, along with each primers sequences and PCR conditions. One group of primers (A:C, A:D, B:C, B:D) will generate products if this insertion is present, while a second group (A:E, A:F, B:E, B:F) should generate products if the insertion is absent. Only the second group of PCRs generated products, consistent with the absence of this insertion and a misassembly of the genome. We sequenced the generated PCR products across the ends of DiNV, which confirmed the *Serratia* misassembly, to NCBI (accession: MF966380).

Because considerable viral genetic variation existed within this individual *Drosophila* male, we sought to generate a consensus DiNV sequence. To that end, we called high frequency variants using GATK HaplotypeCaller (parameters: --ploidy 10), which we then inserted into the sequence using GATK FastaAlternateReferenceMaker, resulting in a final circular genome, 155555bp long (DePristo et al., 2011). The genome and annotation is available at NCBI accession number MF966379 (to be released upon acceptance of the manuscript).

### 2.4. DiNV gene identification and content

We identified the gene content of DiNV based on methods used previously (Wang et al., 2012, 2008, 2007; Yang et al., 2014). We predicted methionine-initiated open reading frames (ORFs) encoding 50 amino acids or more and showing minimum overlap using ORF Finder (http://www.ncbi.nlm.nih.gov/gorf/gorf.html) (Rombel et al., 2002), the putative coding regions were numbered as DiNV ORFs. We first used BLASTP and BLASTN to identify orthologs in a database of all nudivirus ORFs (-evalue 0.0001, downloaded from the NCBI gene database in October 2016) and performed reciprocal BLASTP and BLASTN searches versus Kallithea virus, *Oryctes rhinoceros* Nudivirus (OrNV) and *Gryllus bimaculatus* Nudivirus (GrBNV) to confirm the hits found previously. Following this we confirmed each ORFs annotation via BLASTP and BLASTN to the NCBI non-redundant database using default parameters with an e-value cutoff of 0.0001. All ORFs were investigated for characteristic sequence signatures using the conserved domain search tool (http://www.ncbi.nlm.nih.gov/Structure/cdd/wrpsb.cgi) and Pfam with an E-value cutoff of 1 (Finn et al., 2016), with any identified domains recorded in table S2.

### 2.5. DiNV divergence evolution

We identified genes that may be evolving under positive selection between DiNV and its two closest relatives, Kallithea virus and *Oryctes rhinoceros* Nudivirus, by comparing the rates of nonsynonymous to synonymous divergence in each of the 85 shared ORFs. We aligned each set of orthologous nucleotide sequences using PRANK (parameters: -codon +F) (Löytynoja, 2014). Using these codon-based alignments, we found codons shared across all genomes and calculated non-synonymous and synonymous divergence using a custom Biopython script. In this script, we parsed the PRANK generated phylip files for each ORF and identified codons present in both genomes. Using the standard codon table, we identified the number of codons with nucleotide substitutions resulting in an amino acid change (non-synonymous), the number of codons with substitutions resulting in no change (synonymous), and the number of possible synonymous and non-synonymous substitutions for all shared codons in each ORF. For each ORF, we used these numbers to find the proportion of non-synonymous substitutions of all possible non-synonymous substitutions (dN), the proportion of synonymous substitutions of all possible synonymous substitutions (dS), and dN/dS.

Following this we also defined amino acid substitutions as either radical (to an amino acid of a different group based on their side chains) or conservative (to an amino acid with a similar side chain – in the same group) (Smith, 2003). For a broader view of genome-level evolution, we aligned each genome using lastZ to identify blocks of synteny which we visualized using RCircos (Rahmani et al., 2011; Zhang et al., 2013).

We also concatenated the 20 conserved genes from all nudiviruses and AcMNPV, aligned these sequences using MAFFT (Katoh et al., 2002) and generated a phylogeny using PhyML (Guindon et al., 2010) to place DiNV in the nudivirus phylogeny.

### 2.6. DiNV population genetics

Because we found considerable within-host DiNV genetic variation, we identified polymorphisms in ICH01M DiNV. For this we used Lofreq (Wilm et al., 2012) and allowed for the detection of indels (Lofreq parameters: indelqual –dindel, call –call-indels –min-mq 20), we considered polymorphisms with a minimum frequency threshold of 0.002, which corresponds to about two-fold coverage of a specific site (Wilm et al., 2012). We also filtered these SNPs for polymorphisms exclusively at synonymous sites.

Using all variation detected with Lofreq (and synonymous variation), we performed a genome wide scan of within host polymorphism to find Watterson's theta, Tajima's pi and Tajima's D across sliding windows and within each gene, using Popoolation (Kofler et al., 2011; Tajima, 1989). We also performed McDonald-Kreitman tests (McDonald and Kreitman, 1991) with either Kallithea virus or OrNV as the outgroup and calculated alpha (the proportion of adaptive substitutions) (Smith and Eyre-Walker, 2002) between each genome and DiNV using a custom Biopython script and the gene codon alignments generated by PRANK previously for the estimation of dN/dS.

We also calculated a simulated neutral expectation of Tajima’s pi and Tajima’s D for the genome based on a population growth model in ms (Hudson, 2002). We estimated this expectation using both the silent and total estimates of Watterson’s theta, the estimated population size from Lofreq (1000) and the median growth rate taken from across a range of viruses (0.48). We then compared our simulated 2.5^th^ and 97.5^th^ quantiles to the observed quantiles for both silent and total polymorphism.

## 3. Results & Discussion

### 3.1. DiNV structure and genes

Following an iterative assembly approach, we found the DiNV genome is 155,555bp, making it among the larger genomes for sequenced nudiviruses (Bézier et al., 2015) and slightly larger than its closest relative, the Kallithea virus (152,390bp) (Webster et al., 2015). The DiNV GC content (30%) is also comparable to other nudiviruses which range from 25 to 42% GC (Bézier et al., 2015). We found 107 ORFs (Figure 1A, Supplementary Figure 1, Supplementary Table 2), resulting in a coding density of 71.7%, similar to Kallithea virus, but on the low end of coding densities for nudiviruses and much lower than all other baculoviruses (Bézier et al., 2015; Wang et al., 2012). DiNV, shares 89 (83%) of its ORFs with the other *Drosophila* nudivirus, Kallithea virus, 85 (79%) ORFs with its next closest relative, OrNV, and 68 (64%) ORFs with GrBNV. Not surprisingly, the 68 ORFs found in all four genomes, include all 20 of the core conserved baculovirus ORFs that are necessary for baculovirus function: ORFs associated with late and very late gene transcription (*P47*, *LEF*-8, *LEF*-9, *LEF*-4, *VLF-* 1, and *LEF*- 5), replication (*DNA polymerase* and *Helicase*), virus structure (*P74, PIF*-1, *PIF-*2, *PIF*-3, *AC68*, *VP91*, *VP39*, *38K*, *PIF-4/19kda* and *ODV-E56*), and those of unknown function (*AC81* and *AC92*) (Jehle et al., 2006; Wang et al., 2012; Wang and Jehle, 2009). Protein identity of these 20 ORFs between DiNV and Kallithea virus ranges from 0.16 to 0.94 (median = 0.75), between DiNV and OrNV ranges from 0.23 to 0.98 (median = 0.83) and between DiNV and GrBNV ranges from 0.35 to 0.99 (median = 0.67).

**Figure 1:**
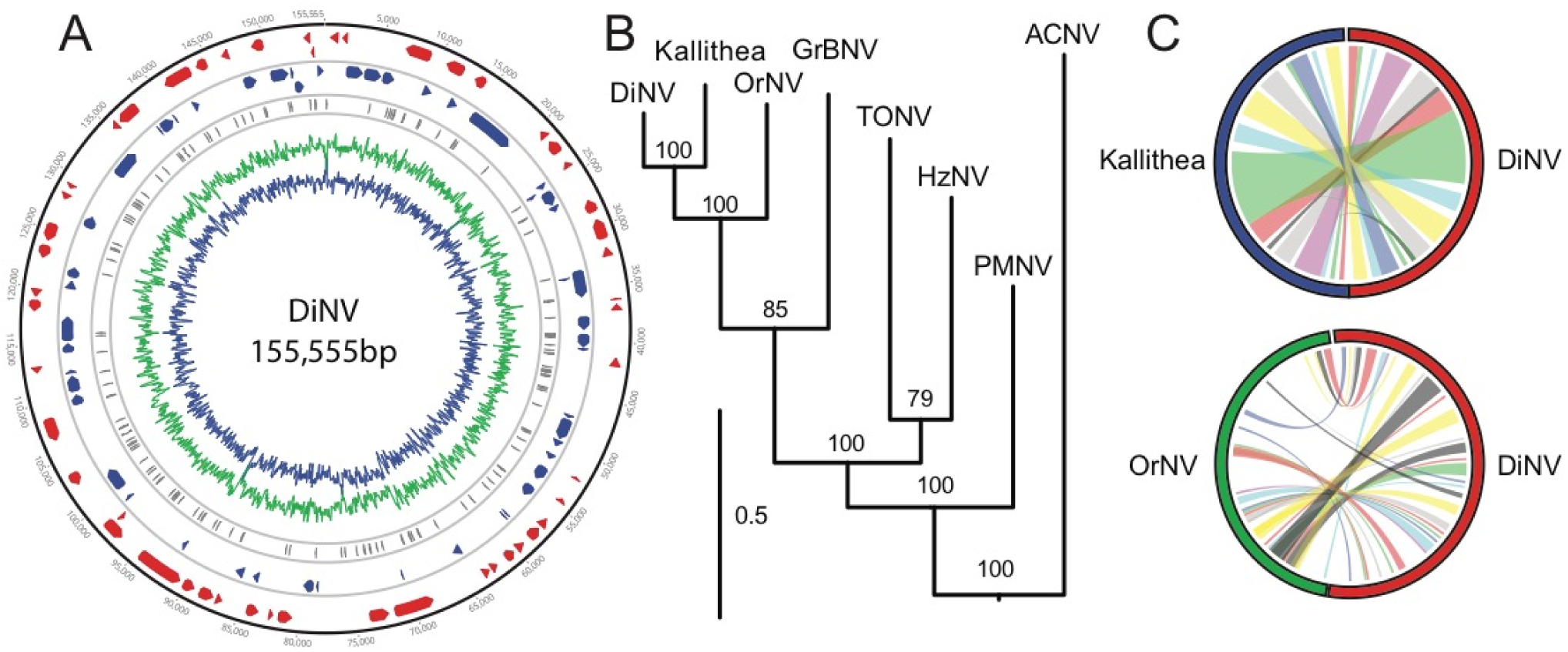
DiNV genome and its relation to other nudiviruses. A) DiNV genome map. The genome is 155555bp, containing 107 ORFs. Anti-sense ORFs are shown in red, Sense ORFs in blue and repeat regions are shown in grey. The percent of AT/GC content is show across the genome in green/blue. B) DiNV on a nudivirus maximum-likelihood phylogeny, using nucleotide sequences of the 20 core ORFs found in all baculoviruses. DiNV is a sister genome to Kallithea with OrNV as its next closest genome. Each branch point shows the bootstrap support from 100 bootstrap replicates, with a scale bar representing 50% divergence. C) DiNV synteny with Kallithea virus and OrNV. Colors are randomly assigned, with extensive blocks of synteny separated by regions with no assignable orthology. Notice that gene order and the size of synteny blocks declines as viruses become more diverged.

Like other annotated nudiviruses, we also find a *polyhedrin/granulin* ORF (*polh/gran*), homologous to the lepidopteran gene. It is unclear what role this gene plays in the nudivirus lifecycle, or its function in its atypical occlusion bodies. Additionally, previous work suggests that the nudivirus *polh/gran* may be the result of convergent evolution, and is unrelated to baculovirus *polh/gran* (Coulibaly et al., 2009, 2007; Wang and Jehle, 2009). It is generally thought that *polh/gran* stabilizes baculovirus virions (Coulibaly et al., 2007; Rohrmann, 2013), so may perform a similar role in the stable formation of virion in nudivruses.

*ODV-E56* appears to be duplicated in both DiNV and Kallithea virus, with a novel copy at 5.5kbp (*ODV-E56-2*) and the original at 122.8kbp. A maximum likelihood phylogeny of *ODV-E56* nucleotide sequences from nudiviruses suggests this duplication occurred before the DiNV-Kallithea divergence (Supplementary Figure 2).

**Figure 2:**
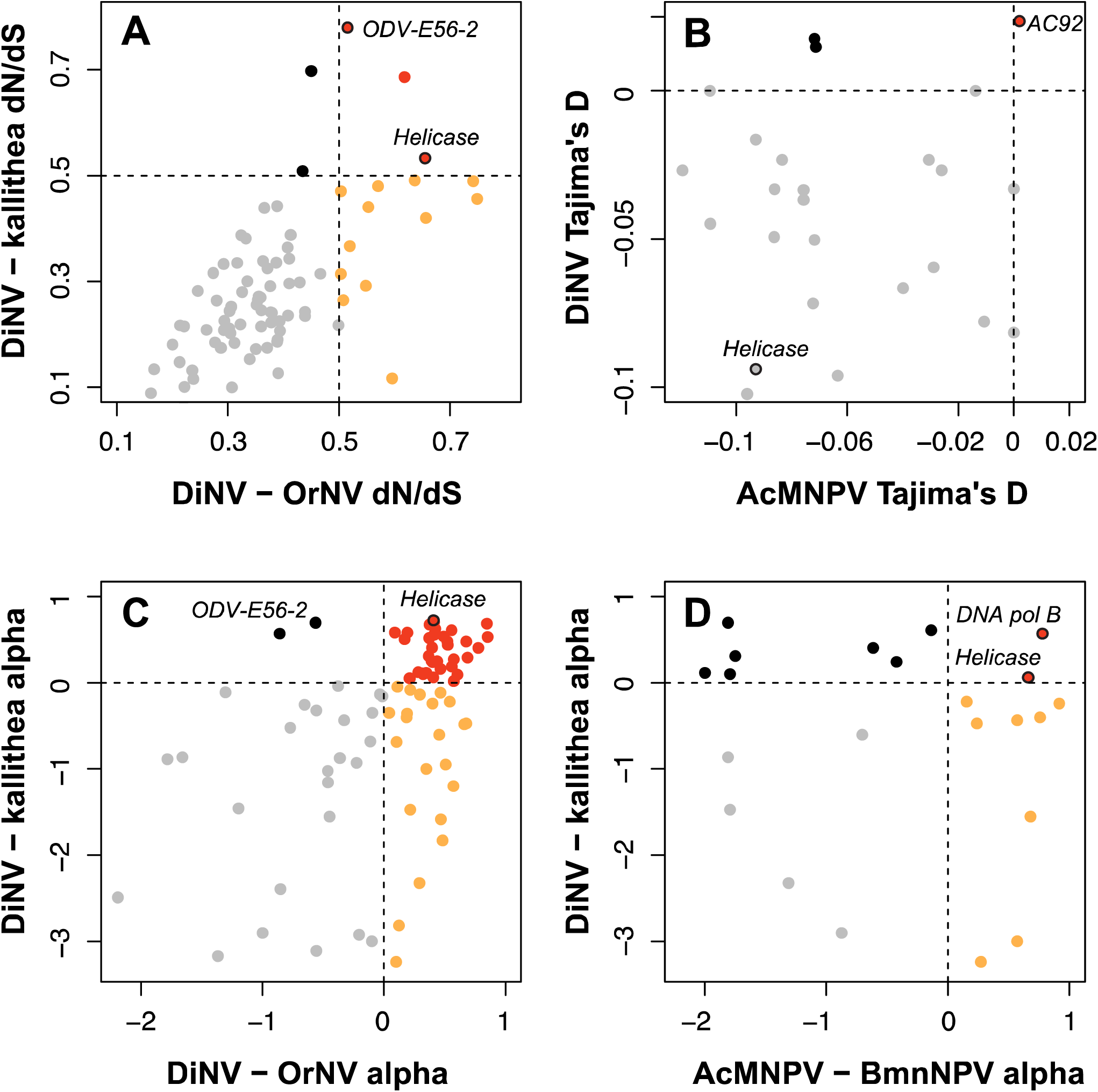
Evolution of DiNV ORFs. For each comparison, we assigned a cut off, either arbitrary to indicate less constrained purifying selection (in the case of dN/dS) or to indicate natural selection (in the case of alpha and Tajima’s D). ORFs above the cutoff in both comparisons are colored red, those above the cutoff in exclusively the OrNV (or AcMNPV/BmNPV) comparison are colored orange, those above the cutoff in exclusively the Kallithea virus comparison are colored black and those below the cut off in both cases are colored grey. A) dN/dS of DiNV ORFs using Kallithea virus and OrNV as paired sequences with an arbitrary cutoff of 0.5 shown (dotted line). Very few genes show adaptive evolution in both comparisons. B) Tajima’s D (a measure of selection within a population) for ORFs shared between AcMNPV and DiNV. C) Alpha (the proportion of adaptive amino acid substitutions, from Mcdonald Kreitman tests) between DiNV – Kallithea and DiNV – OrNV. D) Alpha compared for AcMNPV and DiNV. Only two genes overlap with adaptive substitutions, Helicase and DNA polymerase.

The 16 putative ORFs unique to DiNV show no significant difference in GC-content or length from previous described proteins (Mann-Whitney U Test p-value = 0.87, W = 1375). Among these 16 novel ORFs, 6 have motifs shared with other proteins including a thymidylate synthase, a maturase domain for intron splicing, a T-cell activation factor, a glycosylation protein, a transcription factor domain and a Gastropod egg laying hormone precursor protein domain (Supplementary Table 2).

The genome is comprised of 5.1% simple repeats dispersed across 156 regions (Figure 1A in grey, Supplementary Table 2). These repeats are primarily AT-rich (e.g. ATAT, ATTT, TAATTA, TTGATA), contributing to the low GC content seen throughout the genome (the genome is 33.9% GC after removing repeats). When comparing the densities of repeats within and outside of coding regions, we find no significant difference in the density of repeats between regions (Mann Whitney U test W = 1138, p-value 0.9476), and no excess of repeats in the larger non-coding regions (> 1000bp) versus the smaller regions (< 1000bp) (Mann Whitney U test W = 1297, p-value 0.2067).

A nudivirus phylogeny built using the amino acid sequences of the 20 ORFs shared across all baculoviruses (Figure 1B), shows that DiNV clusters with Kallithea virus and OrNV, confirming previous results with fewer loci (Unckless, 2011). Most of the DiNV genome is syntenic with Kallithea virus, with slight differences in gene content and position of ORFs (Figure 1C, Table S2). However, DiNV shows much less gene retention or synteny with OrNV (Figure 1C, Table S2) and we were unable to find regions of synteny for blocks larger than individual genes for more divergent nudiviruses including GrBNV (Table S2). These results are consistent with other nudivirus and baculovirus studies which found both gene content and synteny are poorly conserved (Wang et al., 2012).

### 3.2. Nudivirus evolution within and between hosts

We suspect that positive selection observed between OrNV and the two *Drosophila* infecting nudiviruses may be due to adaptation to a new host system. To test this, we looked for signatures of adaptation between genes DiNV shares with both Kallithea virus and OrNV.

We calculated dN/dS, the proportion of non-synonymous substitutions to synonymous substitutions between DiNV and Kallithea virus, and DiNV and OrNV. Most proteins are under purifying selection in both cases (dN/dS < 1), with no ORFs, in either comparison, showing evidence of strong positive selection, suggested by a dN/dS greater than 1. The functional category with the median highest dN/dS are involved in host infection (e.g. *VLF-1*, *PIF-1*, *PIF-3*), suggesting that these genes may be important to adapting to a new host, though this group is not a statistical outlier (81^st^ percentile based on permutations). As we find no signatures of positive selection, we attempted to identify genes under unconstrained evolution or putative adaptation and identify genes which overlap in several analyses looking for adaptation, hoping to infer which genes are the most likely to be undergoing adaptation within and between hosts. Using an arbitrary threshold of dN/dS > 0.5 for unconstrained evolution/putative selection, *Helicase*, *ODV*-*E56-2* and a hypothetical protein are the only ORFs to not show signatures of purifying selection in both comparisons (Figure 2A). These results with *Helicase* are consistent with previous findings which show *Helicase* is one of the most rapidly evolving genes across baculoviruses and nudiviruses (Hill and Unckless, 2017). *Helicase* has previously been strongly implicated in host range expansion of baculoviruses (Argaud et al., 1998; Croizier et al., 1994), so unconstrained evolution of viruses across differing host species is not unexpected. Only two hypothetical ORFs are above the 0.5 threshold exclusively in the DiNV/Kallithea virus divergence (Figure 2A, black points). Interestingly, twelve genes have dN/dS above 0.5 exclusively between DiNV and OrNV (Figure 2A, orange points). These twelve include *LEF-3*, *GrBNV_gp28*-*like* protein, and ten other hypothetical proteins, including two trypsin-serine proteases and one patatin phospholipase. As expected, most ORFs are under purifying selection, likely because they are close to a fitness optimum, with few changes being adaptive. While Kallithea virus and DiNV are found in similar hosts, OrNV infects a strikingly different host organism, *Oryctes rhinoceros*. Thus, the higher rate of amino acid substitutions in these ORFs between DiNV and OrNV may be important for adaptation to a new host system.

Using the divergence data between DiNV and Kallithea virus or OrNV coupled with polymorphism in DiNV, we calculated the proportion of adaptive substitutions in each gene (alpha) using the McDonald-Kreitman test (McDonald and Kreitman, 1991). This was done using polymorphism found in the virus found in a single host, so it may not necessarily represent the entire population. Values of alpha greater than zero indicate that some amino acid substitutions were fixed by natural selection in that gene (Smith and Eyre-Walker, 2002).

Fifty-nine of the ORFs have an alpha value greater than zero, thirty-five in both comparisons, suggesting these genes have at least one substitution fixed by natural selection (Table S2). None of these genes have a significant result from a McDonald-Kreitman test, likely due to the small number of polymorphic sites (Chi-squared test p-value > 0.23). Eight of the ORFs with putatively adaptive substitutions are among the core 20 baculovirus ORFs (*Helicase*, *19K*, *DNA polymerase*, *P74*, *polyhedrin/granulin*, *VLF-*1, *Ac92* and *PIF*-3), as well as a ligase and 26 hypothetical proteins (Figure 2C). Consistent with the divergence analysis, we find two only two ORFs (*ODV-*E56-2 and a hypothetical protein) with adaptive substitutions exclusively between DiNV and kallithea virus, versus 22 ORFs (*P47*, *VP39*, *VP91*, *PIF*-1, *PIF*-2, *LEF*-3, *LEF*-4, *ribosomal reductase* 1, *ribosomal reductase* 2, *61K*, *AC81* and 11 hypothetical proteins) with adaptive substitutions between DiNV and OrNV. Among the 20 core baculovirus ORFs, only *Helicase* has adaptive substitutions in all tests. This is also consistent when looking across baculoviruses in general (Hill and Unckless, 2017). A similar analysis was performed on a relatively closely related baculovirus, AcMNPV, comparing the results of these two surveys, we find that *Helicase* and *DNA polymerase* are the two ORFs with alpha greater than 0 for both the DiNV and AcMNPV analyses (Figure 2D, Supplementary Table 2) (Hill and Unckless, 2017). *Helicase* has previously been implicated in the extension of host range for a baculovirus (Argaud et al., 1998; Croizier et al., 1994), so putatively selected changes between host species comes as no surprise, however unconstrained changes between similar host species is surprising.

Most specific amino acid changes between DiNV and OrNV are either to aliphatic or uncharged residues (3592 and 3003 respectively, of 10734 changes), a similar proportion to the standing amino acid types (11035 and 11501 respectively, of 37833 amino acids). One sign that natural selection is driving sequence divergence is if amino acid changes are more likely to be ‘radical’ changes than expected by chance e.g. changing to a different amino acid type (polar-uncharged, polar-acidic, polar-basic, non-polar-aliphatic, non-polar-aromatic and other non-polar). A significant proportion of changes are radical compared to ‘conservative’ changes to similar amino acids (Wilcoxon paired test: W = 40213, p-value = 1.31e-11). However, when categorizing the data by ORF functional group (e.g. replication, transcription, host-infection) or individual ORF, we find no significant excess of radical changes in any ORFs (Wilcoxon paired test p-value > 0.21), with no effect of functional category (*p*- value > 0.12). Polymorphic amino acid changes seen in the virus are also primarily to aliphatic or uncharged amino acids from any amino acid type, with no difference in the ratio of conserved to radical changes seen at any level (Wilcoxon paired test W < 191 p-value > 0.32).

#### Evolution within DiNV

Recent adaptive evolution is characterized by reduced DNA polymorphism in the region surrounding the selected locus and an excess of rare mutations compared to the neutral expectation. The Tajima's D statistic allows for the detection of this: a negative Tajima's D is consistent with a recent selection at an ORF due to an excess of low frequency derived polymorphism, while a positive Tajima’s D suggests balancing selection and maintained polymorphism (Tajima, 1989). We calculated the per site Tajima's D both using a sliding window approach across the genome of DiNV and by individual ORFs, using SNPs called from the pool of DiNV particles infecting a single individual, ICH01M. Given the evidence for recombination in related viruses (Hill and Unckless, 2017; Rohrmann, 2013), natural selection can leave signatures in specific regions of the genome.

Tajima's D is mostly negative across the viral genome (78 ORFs have Tajima’s D < 0), consistent with the fact that the viral population size is much reduced upon initial infection, then increases as the infection proceeds. We simulated the expected Tajima’s D in a population growth model using *ms* (Hudson, 2002), and no ORFs were below the 2.5^th^ quantile of the simulated distribution (-0.149), suggesting no deviation from the neutral expectation, similar to our dN/dS results. Because the detection of sweeps may be affected by the action of direct selection on non-synonymous polymorphism, we also estimated Tajima’s D again using only synonymous sites. Again, we find no ORFs are below the 2.5^th^ quantile of the simulated expectation of Tajima’s D (-0.153).

Though Tajima’s D does not differ from the simulated expectation, we find that Tajima’s D is mostly negative, and varies across the genome, consistent with differing signatures of selection across the genome. We consider regions in the lower 2.5 percentile of Tajima’s D to be the most likely to have recently undergone selection (Figure 2B, Supplementary Figure 3). These windows include only 5 genes: 2 hypothetical proteins, *ODV-E56-2*, *Helicase* and *61K*. These ORFs are also in windows below the 2.5th percentile for pairwise diversity (Supplementary Figure 3, Supplementary Table 2). When analyzing only synonymous sites, in windows below the 2.5^th^ empirical percentile for observed synonymous Tajima’s D, we only find one ORF, ORF59, a trypsin-serine protease not found in the previous survey (Figure S3, Table S2).

*Helicase* is involved in the replication of viral DNA by its unwinding, and is found in a strongly conserved gene cluster found in all baculoviruses (Herniou et al., 2003; Hill and Unckless, 2017; Rohrmann, 2013; Wang et al., 2012). Our results suggest that *Helicase* may be a common target for host suppression, as it contains a conserved domain and is vital to viral replication. This may explain *Helicase*s frequent signatures of unconstrained evolution, positive selection and selective sweeps, as alleles that evolve to escape this suppression are positively selected, resulting in the signatures we observe here (Hill and Unckless, 2017). In fact, previous genetic mapping has found that variation in host range, and ability for host swapping is primarily due to sequence variation in the *Helicase* sequence (Argaud et al., 1998; Croizier et al., 1994; Miller and Albert Lu, 1997). While *Helicase* frequently shows signatures of adaptation across baculoviruses (Hill and Unckless, 2017), thus far, *ODV-E56* shows strong signatures of selection in only the *Drosophila*-infecting nudiviruses (the duplicated copy) and in the alphabaculovirus clade (the original copy), a group of viruses limited to closely related lepidoptera hosts. We looked for evidence of gene conversion between both *ODV-E56* copies, which could lead to patterns like signatures of adaptation. Apart from the first site, there is no shared polymorphism between the two copies and no evidence of gene conversion.

In some windows across the genome, high values of Tajima’s D and pairwise diversity suggest that genetic variation is maintained by balancing selection (Figure 2B, Supplementary Figure 3). We find 14 ORFs have Tajima’s D above 0, and 7 above the 97.5^th^ percentile for the simulated estimate of Tajima’s D (0.0527), while only 2 ORFs were in windows above the upper 97.5th percentile of the empirically estimated pairwise diversity and Tajima’s D (*AC92*, *LEF-*9). Using only synonymous polymorphism, we again find two ORFs in windows above the 97.5^th^ percentile for both Tajima’s D (0.08) and pairwise diversity (*AC92* and ORF81, a putative deoxynucleoside kinase). *AC92* was also found to have the highest Tajima’s D and pairwise diversity estimates in a population of AcMNPV (Hill and Unckless, 2017), suggesting that variation may be being maintained in this ORF in several baculoviruses due to some selective mechanism (Figure 2D). *AC92* is a sulfhydryl oxidase that is thought to form a disulphide bond with the host *p53* to interrupt its action (Hakim et al., 2011; Hou et al., 2012; Long et al., 2009) and with the viral ODV for virion formation, though it is uncertain what role this protein plays in baculovirus infection (Rohrmann, 2013). It’s possible that variation is maintained in this ORF due to its involvement in multiple functions, where different substitutions are beneficial for the proteins separate functions.

## 4. Conclusions

The assembly and annotation of the DiNV genome provides the basis for the development of a powerful new model system for the study of host/DNA virus interaction. The structure of the DiNV genome is largely like other nudiviruses but contains a relatively low percent coding content and several regions with repeated arrays. Several of the genes in DiNV that show selective signatures are not only under selection since the transition from an ancestral host to *Drosophila*, but also show signatures of selection in other baculoviruses. This suggests that in baculoviruses and nudiviruses, only a few key genes are consistently evolving in an adaptive arms races with their hosts.

## 5. Acknowledgements

We thank Brittny Smith for help preparing the library for sequencing the infected fly, Todd Schlenke at the University of Arizona for assistance with fly collections, the Southwest Research Station in Portal, Arizona for allowing us to collect *Drosophila* and two anonymous reviewers for their helpful comments and recommendations for the presented work. Work for this grant including next generation sequencing at the University of Kansas Genome Sequencing Core Laboratory and other supplies and travel was supported in part by the University of Kansas Center for Molecular Analysis of Disease Pathways (NIH P20 GM103638). This project was supported by a Max Kade Postdoctoral Fellowship to TH and NIH grant 4R00GM114714-02 to RLU.

## Supplementary Materials

**Supplementary Table 1:** Primers designed for P47 and LEF-4 based on previous sequencing data (Unckless, 2011).

**Supplementary Table 2:** Table of genes, satellites, positions, closest orthologs for OrNV, GrBNV & Kallithea, Mcdonald Kreitman alpha values, Tajima’s D values, and dN/dS values.

**Supplementary Data 1:** DiNV genome in fasta format

**Supplementary Figure 1:** Image of DiNV genome showing ORFs, satellites and AT/GC content.

**Supplementary Figure 2:** Maximum-likelihood phylogeny of *ODV-E56* genes including the duplications. Bootstrap support values from 100 replicates are shown on the nodes, and a scale bar shown.

**Supplementary Figure 3:** Signatures of positive selection in the DiNV genome. Pairwise diversity (pi) and Tajima's D calculated across the genome in 2kb windows, with 500bp step sizes for total polymorphism and synonymous polymorphism. Values for specific ORFs were confirmed to be in the ranges of the windows by calculating Tajima's D per ORFs. Positions above and below the dashed line are above and below the upper and lower 2.5^th^ percentile of Tajima's D and pairwise diversity in windows along the genome. Positions marked with a cross are ORFs with the entire sequence above or below the percentile cutoffs.

